# Evaluation of antibiotic and peptide vaccine strategies for mirror bacterial infections

**DOI:** 10.1101/2025.10.10.681688

**Authors:** Alexander Kleinman, Joe Torres, Brian Wang

## Abstract

Recent advances in synthetic biology have raised concerns that the creation of bacteria composed completely of mirror-image biomolecules may be possible in the coming decades. Such “mirror bacteria” could pose an unprecedented biosecurity risk, as they are predicted to be highly virulent pathogens resulting from severe immune evasion. Pharmaceutical interventions, including antibiotics and vaccines, could likely provide some limited protection in the event of a mirror bacterial outbreak. Understanding the feasibility of specific pharmaceutical antimirror strategies may help inform the appropriate measures to confront the risks of mirror bacteria. Here, we experimentally evaluated the prospects of previously proposed antimirror antibiotic and vaccine strategies. First, we assessed the expected efficacy of existing antibiotics chloramphenicol, linezolid, tedizolid, and aztreonam against mirror bacteria by characterizing the antibacterial activities of their enantiomers against natural-chirality bacteria. We found that *ent*-chloramphenicol, *ent*-linezolid, *ent*-tedizolid, and *ent*-aztreonam exhibited minimal antibiotic activity, suggesting that their parent antibiotics would be ineffective against mirror bacterial infections. Second, to explore whether the enantiomers of existing antibiotics could be suitable as antimirror antibiotics, we evaluated the acute toxicities of *ent*-chloramphenicol, *ent*-linezolid, *ent*-tedizolid, and *ent*-aztreonam in mice, finding that these compounds have favorable acute toxicity profiles compatible with their continued development. Finally, we investigated D-peptide-based approaches to antimirror vaccines, finding that three bacterially-derived D-peptides induced robust D-peptide-specific antibody responses in mice when conjugated to a carrier protein and adjuvanted. Our results support previous suggestions that the enantiomers of existing antibiotics and D-peptide conjugate vaccines represent feasible pharmaceutical strategies against mirror bacterial infections.

**IMPORTANCE:** Technologies enabling the creation of mirror bacteria—bacteria constructed of molecules that are mirror images of those found in nature—may be on the horizon. If created, mirror bacteria would likely pose an unprecedented risk to life on earth given their evasion of natural immune responses. Although they would not protect against widespread environmental damage, antibiotics and vaccines could provide some limited protection to human populations in the event of a mirror bacterial outbreak. We experimentally assessed the prospects of three antimirror strategies: repurposing existing antibiotics, developing mirror versions of existing antibiotics, and developing vaccines based on mirror-image peptides. We broadly conclude that many existing antibiotics are unlikely to be effective against mirror bacteria, while the mirror images of existing antibiotics and mirror-image peptides could serve as foundations for antimirror antibiotics and vaccines, respectively. Our findings may help inform discussions on the proper role of medical countermeasures to defend against mirror bacteria.

## INTRODUCTION

“Mirror bacteria,” or bacteria composed entirely of macromolecules with inverted chirality from their natural counterparts, have recently been recognized as a possible unprecedented risk to human health and the environment (1). In all known bacteria, chiral macromolecules are predominantly built from building blocks with a single chirality (D-saccharides and L-amino acids), and bacteria predominantly employing building blocks of the opposite chirality are not thought to exist naturally (2). However, continued advances in the chemical synthesis of mirror-image biomolecules and in synthetic cell development may jointly enable the creation of synthetic mirror bacteria in the coming decades (3). If created and released into the natural world, mirror bacteria may grow relatively unchecked by natural biotic constraints that rely on chiral interactions (e.g., phage infection, protist predation, and much of bacterial warfare with antimicrobial toxins), promoting their persistence in natural environments and their consequent routine access to multicellular hosts (4). Mirror bacterial infection of these hosts would be expected to be accompanied by severe evasion of natural immune defenses due to defects in mirror biomolecule recognition, processing, and presentation as well as resistance to many immune effector mechanisms (e.g., peptidoglycan cleavage and phagocytosis) (5). Humans with immunodeficiencies that approximate these anticipated immune defects even partially—such as IRAK-4/MyD88, C1/C2/C4, and MHC class II deficiencies resulting in impaired pattern recognition, complement activation, and antigen presentation, respectively—typically require intensive prophylactic pharmaceutical intervention to limit life-threatening bacterial infection, although clinical outcomes are often still poor (5). As a result, albeit expected to be incomplete, the protection of humans and other multicellular organisms against lethal mirror bacterial infection might similarly require the use of medical countermeasures, including antibiotics and vaccines.

Some possible strategies for the development of antibiotics and vaccines against mirror bacterial infections have been previously outlined (6). For example, existing achiral or racemic antibiotics would be expected to straightforwardly maintain their antibiotic activity against mirror bacterial infections. In addition, due to chiral symmetry between enantiomers, the enantiomers of existing chiral antibiotics should exhibit as potent antibacterial activity against mirror bacteria as those existing antibiotics do against the same respective natural-chirality bacteria. Finally, while mirror bacterial antigens (e.g., L-glycans or D-peptides) might only be weakly immunogenic in isolation due to impaired processing and presentation to T cells, conjugation of these mirror bacterial antigens to immunogenic carrier proteins should enable the stimulation of T cell-dependent immune responses (7–9). The enantiomers of existing antibiotics and mirror bacterial antigen conjugates therefore represent possible avenues for antimirror antibiotic and vaccine development, respectively.

However, much remains unknown about the prospects of these strategies for antimirror antibiotic and vaccine development and their comparison to potential alternatives. First, the extent to which existing chiral antibiotics could combat mirror bacterial infections remains uncertain, as not all antibiotics may be equally sensitive to a reversal in chirality. For example, both enantiomers of the quinolone compound merafloxacin demonstrate similar antibacterial activity, whereas for the related quinolone ofloxacin, the (–)-enantiomer is up to 128-fold more potent than the (+)-enantiomer (10, 11). Supporting this variability in chiral sensitivity, computational studies using docking and molecular-dynamics simulations revealed that amoxicillin binds unstably to the mirror-image form of its natural target, Staphylococcus aureus penicillin-binding protein 3, while its enantiomer forms a stable complex with the mirror protein (12). Chiral antibiotics whose enantiomers possess comparable bioactivity against natural-chirality bacteria would have increased promise as antimirror antibiotics, and their existence would have implications for the scope of antibiotic development required to address the threat of mirror bacteria. However, to date, the antibacterial activities of the enantiomers of existing chiral antibiotics remains sparsely characterized, leaving the expected efficacy of these antibiotics against mirror bacteria largely uncertain.

Second, while there would be little uncertainty about the antimirror antibiotic activities of the enantiomers of existing antibiotics, their pharmacokinetic and toxicity profiles may differ from the profiles of their enantiomers and would require independent evaluation. However, so far these profiles have largely not been established.

Lastly, the optimal strategies for enhancing the immunogenicity of mirror bacterial antigens are unclear. Previous studies have indicated that D-peptides conjugated to carrier proteins can induce high levels of anti-D-peptide antibodies, but these studies have typically focused on retro-inverso peptides (D-peptides that possess reversed primary sequences from L-peptides of interest, and therefore that maintain the relative side chain orientations of those L-peptides) (13–17). The observed immunogenicity of retro-inverso peptide conjugates may not represent that of D-peptide conjugates as a whole, given that retro-inverso peptides are designed to mimic immunogenic L-peptides. Additionally, D-peptides formulated with strong adjuvant have been previously reported to elicit T-cell-dependent immune responses comparable to those achievable with L-peptides, providing an alternative potential means of developing D-peptide-based vaccines not reliant on conjugation (18). A dedicated comparison of the immunogenicity of mirror bacterial antigens in combination with conjugation and/or adjuvantation would help to identify the most promising approaches for mirror bacterial vaccine development.

A more detailed understanding of the strategies for antimirror antibiotic and vaccine development would enhance broader discussions around the appropriate role of medical countermeasures in preparing for risks from mirror bacteria. To that end, here we describe a series of initial investigations into these strategies. First, to address the possibility that existing chiral antibiotics might maintain antibacterial activity against mirror bacteria, we selected four approved antibiotics (chloramphenicol, linezolid, tedizolid, and aztreonam) and measured the antibiotic activities of their enantiomers against a diverse panel of bacteria. Second, to preliminarily evaluate the suitability of the enantiomers of existing antibiotics as antimirror antibiotics, we characterized the acute toxicity profiles of *ent*-chloramphenicol, *ent*-linezolid, *ent*-tedizolid, and *ent*-aztreonam in mice. Finally, to investigate strategies for composing antimirror vaccines, we assessed how carrier protein conjugation and adjuvantation affected the immunogenicity of three bacterially-derived D-peptides.

## RESULTS

### Antibiotics chloramphenicol, linezolid, tedizolid, and aztreonam are not expected to maintain their antibacterial activity against mirror bacteria

Over 250 antibiotics are in use in humans globally, of which only a minority are achiral or racemic (6, 19). We therefore set out to understand the extent to which chiral antibiotics, making up the vast majority of our antibiotic armamentarium, might also retain antibiotic activity against mirror bacteria. It is possible to study this in the absence of mirror bacteria by measuring the antibiotic activity of the enantiomers of the antibiotics against natural-chirality bacteria. Thus, we identified four antibiotics whose enantiomers were relatively synthetically accessible—chloramphenicol, linezolid, tedizolid, and aztreonam—for further study. Chloramphenicol is a nitrobenzene compound that inhibits bacterial protein elongation by binding to the peptidyl transferase center of the 50S ribosomal subunit and exhibits broad-spectrum activity against both gram-negative and gram-positive bacteria. Linezolid and tedizolid are first- and second-generation oxazolidinone antibiotics, respectively, that inhibit bacterial protein synthesis initiation by binding to the 23S rRNA of the 50S ribosomal subunit and typically exhibit activity against gram-positive bacteria. Aztreonam is a monobactam antibiotic that inhibits bacterial cell well biosynthesis by binding to bacterial penicillin-binding proteins and exhibits activity against aerobic gram-negative bacteria. Together, these four antibiotics span multiple structural classes, bacterial targets, mechanisms of action, and spectrums of activity.

*ent*-Chloramphenicol, *ent*-linezolid, *ent*-tedizolid, and *ent*-aztreonam were synthesized and their minimum inhibitory concentrations (MICs) were measured against a diverse panel of 18 bacteria, including gram-positive, gram-negative, obligate anaerobic, facultative anerobic, and microaerophilic reference strains or clinical isolates (Table S1). For direct comparison, we also included chloramphenicol, linezolid, tedizolid, and aztreonam. As expected, chloramphenicol demonstrated activity against most bacteria tested, linezolid and tedizolid exhibited strongest activity against gram-positive bacteria, and aztreonam exhibited strongest activity against gram-negative facultative anaerobes (Table 1). Against established quality control organisms, the MIC values of the parent antibiotics typically fell within previously reported reference ranges (20). *ent*-Chloramphenicol, *ent*-linezolid, and *ent*-tedizolid lacked antibacterial activity against any tested bacteria at concentrations up to 128 μg/mL. *ent*-Aztreonam exhibited slight antibacterial activity against only a small subset of bacteria tested, achieving a MIC of 128 μg/mL against *Escherichia coli* and 32 μg/mL against an aztreonam-susceptible isolate of *Enterobacter cloacae*, *Klebsiella pneumoniae*, *Salmonella enterica*, and *Shigella sonnei*. In each of these cases, *ent*-aztreonam was 512- or 1024-fold less potent than aztreonam. These results demonstrate that the antibacterial activities of chloramphenicol, linezolid, tedizolid, and aztreonam are broadly sensitive to a reversal of chirality, and that these antibiotics are not expected to exhibit significant antibacterial activity against a similar panel of mirror bacteria.

**Table 1.** MIC values of chloramphenicol, linezolid, tedizolid, aztreonam, and their enantiomers against a panel of bacteria. Cell colors range from yellow to brown, representing low to high MIC values, respectively. Reference ranges for quality control organisms are included in parentheses. Source numbers, Gram classifications, and oxygen requirements for all strains are provided in Table S1. CHL, chloramphenicol; LZD, linezolid; TZD, tedizolid; ATM, aztreonam; ATM-S, aztreonam-susceptible.

|  | Species | MIC (µg/mL) |  |  |  |  |  |  |  |
| --- | --- | --- | --- | --- | --- | --- | --- | --- | --- |
|  |  | CHL | ent-CHL | LZD | ent-LZD | TZD | ent-TZD | ATM | ent-ATM |
| Gram-negative | *Bacteroides fragilis* | 4 | >128 | 8 | >128 | 1 | >128 | 32 | >128 |
|  | *Bacteroides thetaiotaomicron* | 8 | >128 | 16 | >128 | 1 | >128 | 128 | >128 |
|  | *Campylobacter jejuni* | 4 | >128 | 32 | >128 | 8 | >128 | 16 | >128 |
|  | *Enterobacter cloacae* | 8 | >128 | >128 | >128 | >128 | >128 | 0.5 | >128 |
|  | *Enterobacter cloacae* (ATM-S) | 4 | >128 | >128 | >128 | >128 | >128 | 0.06 | 32 |
|  | *Escherichia coli* | 8 (2–8) | >128 | >128 | >128 | >128 | >128 | 0.12 (0.06–0.25) | 128 |
|  | *Haemophilus influenzae* | 0.5 (0.25–1) | >128 | 32 | >128 | 4 | >128 | 0.12 (0.12–0.5) | >128 |
|  | *Klebsiella pneumoniae* | 8 | >128 | >128 | >128 | >128 | >128 | 0.06 | 32 |
|  | *Pseudomonas aeruginosa* | 128 | >128 | >128 | >128 | >128 | >128 | 4 (2–8) | >128 |
|  | *Salmonella enterica* | 8 | >128 | >128 | >128 | >128 | >128 | 0.06 | 32 |
|  | *Serratia marcescens* | 32 | >128 | >128 | >128 | >128 | >128 | 16 | >128 |
|  | *Shigella sonnei* | 4 | >128 | >128 | >128 | >128 | >128 | 0.03 | 32 |
| Gram-positive | *Clostridioides difficile* | 4 | >128 | 4 | >128 | 0.25 | >128 | >128 | >128 |
|  | *Enterococcus faecalis* | 8 (4–16) | >128 | 4 (1–4) | >128 | 0.5 (0.25–1) | >128 | >128 | >128 |
|  | *Listeria monocytogenes* | 8 | >128 | 8 | >128 | 0.25 | >128 | >128 | >128 |
|  | *Staphylococcus aureus* | 16 (2–16) | >128 | 4 (1–4) | >128 | 0.5 (0.12–1) | >128 | >128 | >128 |
|  | *Streptococcus pneumoniae* | 4 (2–8) | >128 | 2 (0.25–2) | >128 | 0.25 (0.12–0.5) | >128 | >128 | >128 |
|  | *Streptococcus pyogenes* | 4 | >128 | 4 | >128 | 0.25 | >128 | >128 | >128 |

### Acute toxicity is not dose-limiting for *ent*-chloramphenicol, *ent*-linezolid, *ent*-tedizolid, and *ent*-aztreonam in mice

Beyond antibacterial activity, the safety profiles of these antibiotic enantiomers remain unknown but are critical for clinical translation. We investigated the single-dose acute toxicity of *ent*-chloramphenicol, *ent*-linezolid, and *ent*-tedizolid via oral gavage and *ent*-aztreonam intravenously (i.v.) at a high and a low dose in mice for 14 days. For comparison, separate groups of mice were treated with doses of the respective parent antibiotics, which were expected to be well-tolerated. No mortality was observed in animals receiving *ent*-tedizolid, *ent*-aztreonam, or the parent antibiotics (Table 2). However, complete mortality was observed within four hours in groups receiving high doses (2000 mg/kg) of *ent*-chloramphenicol and *ent*-linezolid, while no mortality was observed at the low dose (300 mg/kg).

**Table 2.**
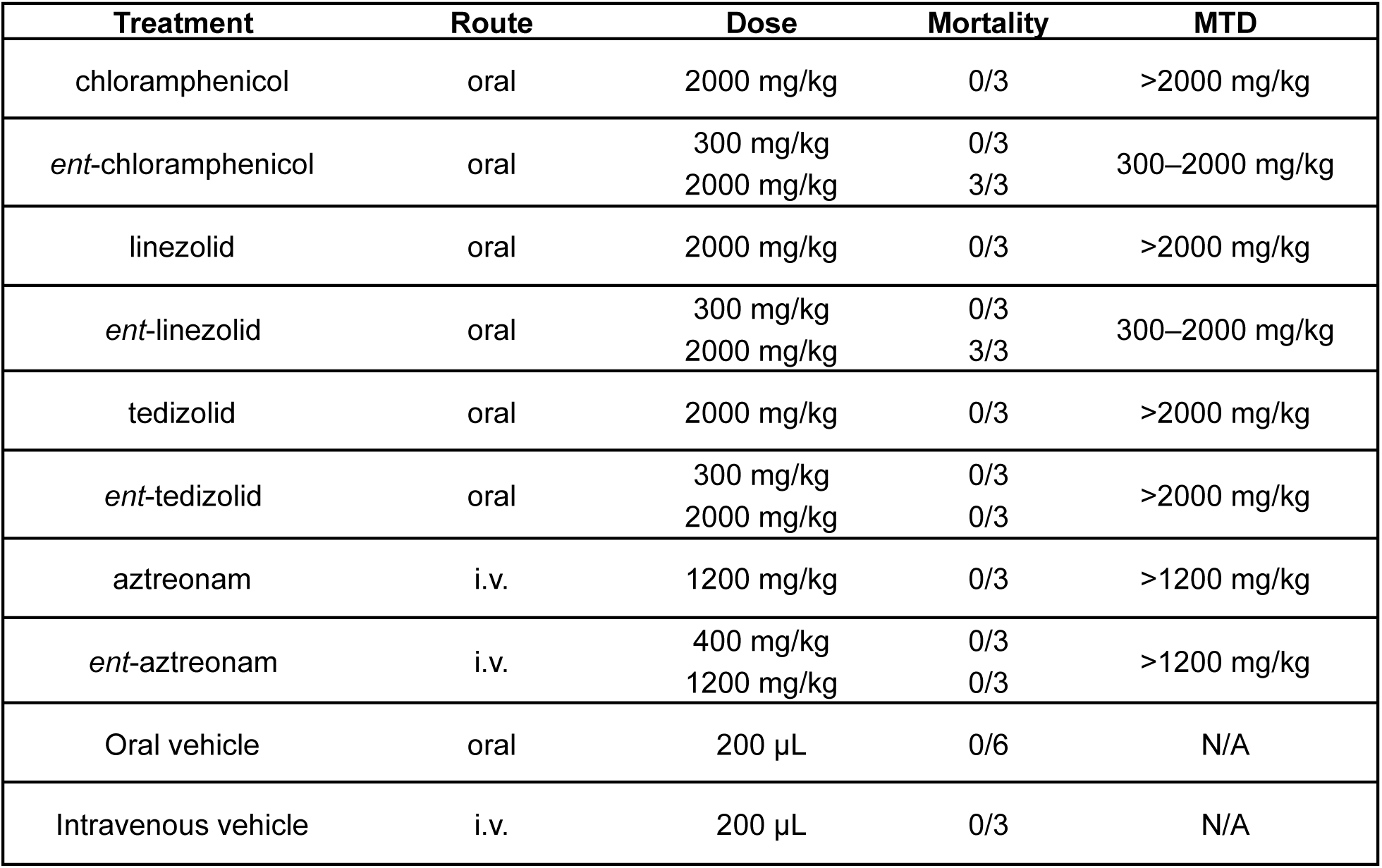
Acute toxicity study design, observed mortality rates, and determined MTDs for *ent*- and parent chloramphenicol, -linezolid, -tedizolid, and -aztreonam.

| Treatment | Route | Dose | Mortality | MTD |
| --- | --- | --- | --- | --- |
| chloramphenicol | oral | 2000 mg/kg | 0/3 | >2000 mg/kg |
| *ent*-chloramphenicol | oral | 300 mg/kg | 0/3 | 300–2000 mg/kg |
|  |  | 2000 mg/kg | 3/3 |  |
| linezolid | oral | 2000 mg/kg | 0/3 | >2000 mg/kg |
| *ent*-linezolid | oral | 300 mg/kg | 0/3 | 300–2000 mg/kg |
|  |  | 2000 mg/kg | 3/3 |  |
| tedizolid | oral | 2000 mg/kg | 0/3 | >2000 mg/kg |
| *ent*-tedizolid | oral | 300 mg/kg | 0/3 | >2000 mg/kg |
|  |  | 2000 mg/kg | 0/3 |  |
| aztreonam | i.v. | 1200 mg/kg | 0/3 | >1200 mg/kg |
| *ent*-aztreonam | i.v. | 400 mg/kg | 0/3 | >1200 mg/kg |
|  |  | 1200 mg/kg | 0/3 |  |
| Oral vehicle | oral | 200 µL | 0/6 | N/A |
| Intravenous vehicle | i.v. | 200 µL | 0/3 | N/A |

Among surviving animals receiving the *ent*-series of compounds, only mild clinical signs were observed infrequently in a few mice. All observed clinical signs uniformly resolved within one day, and all surviving animals appeared normal at study end (Table S2). Necropsy revealed no gross abnormalities in tissues except for variations in uterine thickness and fluid content, consistent with normal estrous cycle-related changes, and white spots on the hearts of two mice receiving *ent*-tedizolid, possibly indicative of dystrophic cardiac calcinosis common in BALB/c mice (21).

All groups receiving the *ent*-series of compounds gained body weight over the course of the study, and no significant differences in body weight change were observed between treatment and control groups on day 14. Among groups receiving the parent antibiotics, only animals receiving tedizolid at a high dose (2000 mg/kg) experienced a significant change in body weight, losing an average of 0.3% body weight by day 14 (*p* ≤ 0.001) (Fig. S1).

There were no significant differences in absolute or relative organ weights among surviving animals receiving *ent*- or parent chloramphenicol or -linezolid compared to their respective vehicle controls (Tables S3 and S4). However, absolute and relative uterus weight in mice receiving high dose (2000 mg/kg) tedizolid and relative liver weight in mice receiving high dose (2000 mg/kg) *ent*-tedizolid were each significantly decreased compared to vehicle controls (*p* ≤ 0.05, Table S5). Additionally, the absolute thymus weight in mice receiving low dose (400 mg/kg) aztreonam was significantly increased compared to vehicle controls, but the same was not true for mice receiving high dose (1200 mg/kg) aztreonam; as a result, this difference was not considered toxicologically meaningful (*p* ≤ 0.05, Table S6).

There were also no significant differences in hematological parameters among surviving animals receiving *ent*- or parent chloramphenicol, -linezolid, or -aztreonam or their respective vehicle controls (Tables S7, S8, and S10). However, mice receiving high doses (2000 mg/kg) of *ent*- and parent tedizolid had significantly decreased hematocrit percentage and significantly decreased mean corpuscular volume, respectively (*p* ≤ 0.05, Table S9).

On the basis of these combined observations, we defined ranges for the maximum tolerated dose (MTD) in mice for each compound (Table 2). The MTD for each of the parent antibiotics exceeds the high doses used in this study (2000 mg/kg for chloramphenicol, linezolid, and tedizolid, and 1200 mg/kg for aztreonam). For *ent*-chloramphenicol and -linezolid, the MTD falls between the low and high doses (300–2000 mg/kg), whereas for *ent*-tedizolid and -aztreonam, the MTD exceeds the high dose (2000 mg/kg and 1200 mg/kg, respectively). Overall, we do not find evidence of severely dose-limiting acute toxicity for *ent*-chloramphenicol, *ent*-linezolid, *ent*-tedizolid, and *ent*-aztreonam that would preclude their further development as antimirror antibiotics, and our results suggest dose ranges for future single-dose or repeat-dose toxicity studies to further define their toxicity profiles.

### D-peptides induce robust D-peptide-specific antibody responses when adjuvanted and conjugated to a carrier protein

We evaluated the immunogenicity of three bacterially-derived synthetic D-peptide antigens (D-OmpB_399_, D-StreptInCor, and D-Omp31-TB) and their L-counterparts, the latter of which have previously been studied as immunogens for protection against bacterial infection (22–24), in combination with an adjuvant and/or conjugation to a carrier protein in BALB/c mice. In total, 24 groups of four mice each were immunized with each antigen, with or without adjuvant (complete/incomplete Freund’s adjuvant, CFA/IFA; or CpG oligodeoxynucleotides) and with or without conjugation to a carrier protein (keyhole limpet cyanin, KLH; or bovine serum albumin, BSA), according to the schedules shown in Fig. 1A. An additional three groups of five mice were immunized with phosphate-buffered saline (PBS) alone as a control for each separate schedule. We measured serum IgG binding antibody responses and, for the Omp31-TB groups, bronchoalveolar lavage fluid (BALF) IgA binding antibody responses by enzyme-linked immunosorbent assay (ELISA) 14 days after the final immunization (Figs. 1B–E).

**Figure 1.**
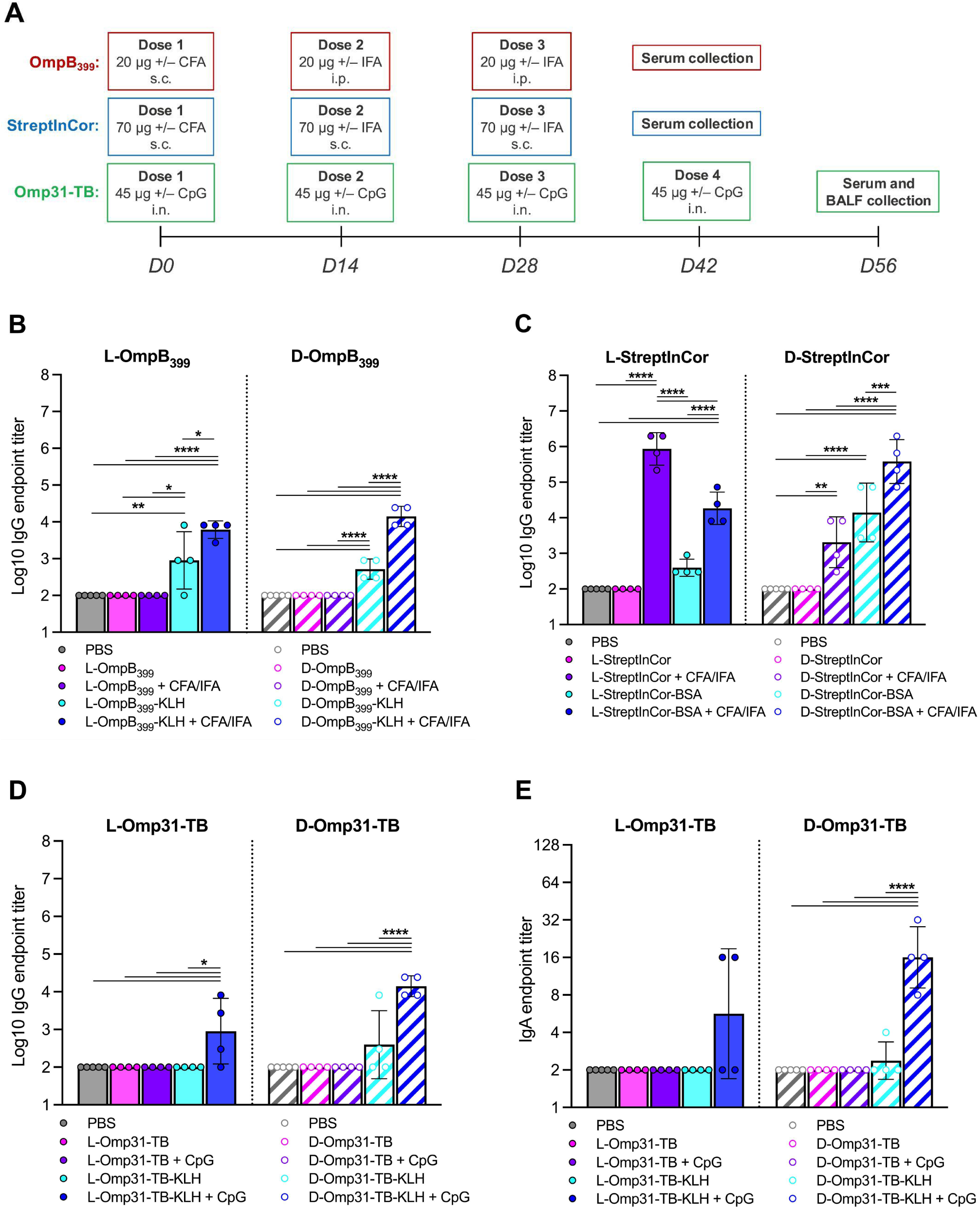
Binding antibody responses to different L- and D-peptide immunization strategies. A) Immunization and collection schedules in BALB/c mice. B–D) Serum IgG antibody levels to B) L- and D-OmpB_399_, C) L- and D-StreptInCor, and D) L- and D-Omp31-TB at 14 days post-final immunization; assay limits of detection were 10^2^. E) BALF IgA antibody levels to L- and D-Omp31-TB at 14 days post-final immunization; assay limit detection was 2. Data is plotted as geometric mean ± geometric SD (*n* = 5, PBS groups; *n* = 4, all other groups). One-way ANOVA with Tukey’s *post-hoc* tests on log-transformed data was used to compare the responses of groups to each peptide. *, *p* ≤ 0.05; **, *p* ≤ 0.01; ***, *p* ≤ 0.001; ****, *p* ≤ 0.0001. s.c., subcutaneous; i.p., intraperitoneal; i.n., intranasal.

Uniformly, L- or D-peptide immunization alone induced no detectable serum antibody responses and no detectable mucosal antibody response for Omp31-TB against the respective peptides, consistent with the weak immunogenicity of peptide antigens generally and D-peptides particularly (25–27). For all three peptides, maximal antibody responses against D-peptides were observed in groups immunized with D-peptide conjugates with adjuvant, followed by groups immunized with D-peptide conjugates without adjuvant. Notably, these robust antibody responses were observed despite the delivery of far fewer peptide instances in groups receiving peptide conjugates, as the amount of antigen was mass-matched across groups and the carrier protein constituted the vast majority of the mass of peptide conjugates. Unconjugated D-peptides with adjuvant induced a moderate level of peptide-specific antibodies for D-StreptInCor, but no detectable serum antibodies against either D-OmpB_399_ or D-Omp31-TB, and no detectable mucosal antibodies against D-Omp31-TB. In general, serum and mucosal antibody levels for D-peptide-immunized groups were comparable to or higher than those of the matched L-peptide-immunized groups, with the exception of groups immunized with StreptInCor plus adjuvant. No mice immunized with L-peptides, with or without adjuvanting or conjugation, produced detectable antibodies against D-peptides (data not shown). Overall, our results indicate that conjugation to a carrier protein robustly enhances the immunogenicity of D-peptides, as with L-peptides; in addition, adjuvantation further augments the immunogenicity of D-peptide conjugates, but adjuvantation alone does not produce the antibody titers achieved with conjugation.

## DISCUSSION

We examined three medical countermeasure strategies against mirror bacteria: repurposing existing chiral antibiotics, developing their enantiomers as antimirror antibiotics, and using adjuvantation and/or conjugation to increase the immunogenicity of D-peptides as possible antimirror vaccines. Our results broadly suggest that existing chiral antibiotics would be ineffective against mirror bacteria, that their enantiomers could be nontoxic and effective antimirror antibiotics, and that adjuvanted D-peptide conjugates may be promising as a basis for mirror bacterial vaccines.

We first evaluated the expected efficacies of approved antibiotics chloramphenicol, linezolid, tedizolid, and aztreonam against mirror bacterial infections by measuring the antibacterial activities of their enantiomers against natural bacteria, finding that none of the enantiomers showed significant antibacterial activity. These data are consistent with previous reports that *ent*-chloramphenicol has minimal antibiotic activity (28, 29), that oxazolidinones typically lose antibiotic activity upon stereochemical inversion at their sole chiral center (30–32), and that the stereochemical configurations of the chiral centers of monobactams are important for their antibiotic activity (33, 34). Notably, chloramphenicol, linezolid, tedizolid, and aztreonam each possess only one or two stereocenters. Most other chiral antibiotics are more stereochemically dense, such that a reversal in the chirality of their targets would be expected to have an even greater detrimental impact on their antibiotic activities. For example, penicillins, cephalosporins, and carbapenems have two to six stereocenters, tetracyclines contain four to six, and macrolides and aminoglycosides possess over ten. The complexity of these antibiotics would also necessitate lengthier total synthesis campaigns for their enantiomers to experimentally evaluate their expected antibiotic activities against mirror bacteria (35). Although we evaluated a limited set of antibiotics, our findings broadly suggest that the straightforward repurposing of existing chiral antibiotics would be ineffective and support the development of dedicated antimirror antibiotics instead.

The enantiomers of existing antibiotics may be promising antimirror antibiotics if they were to have suitable pharmacokinetic and toxicity profiles. Therefore, we next characterized the acute toxicity profiles of *ent*-chloramphenicol, *ent*-linezolid, *ent*-tedizolid, and *ent*-aztreonam in mice, finding that they were generally associated with low acute toxicity as characterized by their MTDs. Human equivalent MTDs calculated using allometric scaling would suggest that *ent*-chloramphenicol and *ent*-linezolid would be maximally tolerated in humans at doses of between approximately 25 mg/kg and 160 mg/kg, and that *ent*-tedizolid and *ent*-aztreonam would be maximally tolerated at over approximately 160 mg/kg and 100 mg/kg, respectively. For comparison, to treat bacterial infections, chloramphenicol, linezolid, and tedizolid are typically prescribed in adults at approximately 3–12.5 mg/kg per dose, and aztreonam is prescribed at approximately 15–30 mg/kg per dose. The calculated human equivalent maximum tolerated doses for the tested *ent*-compounds are thus uniformly higher than the doses currently recommended for their parent antibiotics.

However, we caution against an overinterpretation of these data due to limitations in our study design. First, while we tested each *ent*-compound as their free forms, chloramphenicol and tedizolid are administered clinically as ester prodrugs, which complicates direct dose comparisons. Second, each of the parent antibiotics are dosed repeatedly to treat bacterial infections in human patients. Given that the same would likely be true of the respective *ent*-compounds for treating mirror bacterial infections, repeat-dose toxicity studies are needed to better characterize the relevant toxicity profiles of these *ent*-compounds. Third, additional toxicity studies in non-rodent species would also provide a more detailed understanding of the toxicity profiles of these drugs and are typically recommended by regulatory guidelines (36, 37). Finally, although we may reasonably conclude that the antibacterial activities of the *ent*-compounds against mirror bacteria would be the same as those of the parent antibiotics against natural bacteria, the doses required for clinical therapeutic efficacy may differ substantially. The pharmacokinetic profiles of the enantiomeric compounds may still be distinct, and higher doses of antimirror bacterial drugs may be required to compensate for the severe immune evasion expected to accompany mirror bacterial infections. Any toxicity profile for antimirror antibiotics must therefore be interpreted in light of this uncertainty in dose required for clinical benefit. Nevertheless, our findings on the acute toxicity profiles of *ent*-chloramphenicol, *ent*-linezolid, *ent*-tedizolid, and *ent*-aztreonam support their suitability for further development as antimirror antibiotics, and reinforce the development of enantiomers of existing antibiotics as a medical countermeasure strategy against mirror bacteria.

We finally investigated strategies for the design of mirror bacterial vaccines by examining the immunogenicity of three bacterially-derived D-peptides in combination with adjuvantation and/or conjugation to carrier proteins. Our results suggest that adjuvanted D-peptide conjugates can be robustly immunogenic and may provide a strategy for vaccine development against mirror bacteria, despite the poor immunogenicity of D-peptides in isolation. Our findings corroborate the consistent immunogenicity previously observed with retro-inverso peptide conjugates. Notably, the absolute titers of D-peptide-specific antibodies induced with adjuvanted D-peptide conjugates typically exceeded the absolute titers of L-peptide-specific antibodies induced by the respective adjuvanted L-peptide conjugates. Similarly, retro-inverso peptide conjugates have been observed to induce higher antibody titers than their parent L-peptide conjugates (13, 38). As with retro-inverso peptide conjugates, the source of this enhanced immunogenicity may be the resistance of D-peptides to proteolytic degradation, which prolongs their half-life *in vivo*.

Our results also suggest that longer D-peptides may be suitable as components of conjugate vaccines. The three peptides tested—OmpB_399_, StreptInCor, and Omp31-TB—have lengths of 20, 56, and 16 amino acids, respectively. While OmpB_399_ and Omp31-TB have lengths compatible with direct loading onto MHC class II molecules, StreptInCor requires proteolytic processing for presentation. Given that D-peptides are resistant to such processing, we expected that adjuvantation and/or conjugation would be particularly necessary to boost the immunogenicity of D-StreptInCor. We found that the combination of conjugation and adjuvantation was sufficient to enable D-StreptInCor to be an effective immunogen. Since longer peptides can present conformational epitopes, and antibody responses against conformational epitopes are important for protective efficacy against bacterial infection (39), our findings affirm important avenues for mirror bacterial vaccine design.

Much remains unknown about the potential of D-peptide-based vaccines. The protective efficacy of D-peptide-based vaccines against mirror bacterial infection would be determined by a variety of factors including antibody functionality (e.g., opsonophagocytic capacity), antibody isotype/subclass distribution, and Th1/Th2 bias, each of which may have a complex dependence on the choice of peptide, adjuvant, carrier protein, route of administration, and dose (40–42). The immune markers predictive of protective efficacy would also likely differ between natural and mirror bacterial infections. An understanding of the persistence of the antibody response and the extent to which immunological memory could be established and re-activated would be additionally important to inform vaccine design. Finally, while we have found D-peptide adjuvantation and conjugation to a carrier protein to be associated with higher antibody titers in our studies, prior research on analogous bacterial glycoconjugate vaccines demonstrates that the influence of adjuvantation and conjugation on immunogenicity can be variable, dependent on age and immunization schedule, and not always consistent between animals and humans (43). Future research would be needed to progressively establish general principles for D-peptide-based—or other mirror antigen-based—vaccine design.

Understanding the prospects of strategies for medical countermeasure development is critical for informing a societal response to the threat of mirror bacteria. Our studies highlight the enantiomers of existing antibiotics and adjuvanted D-peptide conjugates as promising directions for antimirror antibiotic and vaccine development, respectively. Our results lay the groundwork for future research on antimirror medical countermeasures, and we hope that this work may productively advance broader discussions around the role of medical countermeasures in confronting the risks of mirror bacteria.

## ACKNOWLEDGEMENTS

This work was supported by funding from Open Philanthropy. We thank the Mirror Biology Dialogues Fund and Dr. John Rex for providing critical reviews of the manuscript prior to submission.

## CReDiT authorship contribution statement

Alexander Kleinman: Conceptualization, Methodology, Investigation, Writing - review & editing. Brian Wang: Conceptualization, Methodology, Formal Analysis, Investigation, Writing - original draft, Writing - review & editing, Visualization, Supervision, Project Administration, Funding Acquisition. Joe Torres: Conceptualization, Methodology, Investigation, Writing - review & editing.

## MATERIALS AND METHODS

### Antibiotics

*ent*-Linezolid, *ent*-tedizolid, and *ent*-chloramphenicol were synthesized by Wuxi AppTec (Shanghai, China). *ent*-Aztreonam was synthesized by Pharmaron (Beijing, China). For *ent*-compounds, structures were confirmed with ^1^H NMR and purities were >95% as verified by LC-MS or HPLC. Linezolid and tedizolid were purchased from MedChemExpress (Monmouth Junction, NJ). Chloramphenicol was purchased from Sigma-Aldrich (St. Louis, MO). Aztreonam was purchased from TCI America (Portland, OR). Levofloxacin (Sigma-Aldrich, St. Louis, MO) and meropenem (United States Pharmacopeia, Rockville, MD) were used as internal assay controls for minimum inhibitory concentration (MIC) testing.

### Bacterial culture and media

Source numbers for all strains used are provided in Table S1. All bacterial culture and testing was performed by Microbiologics, Inc. (Kalamazoo, MI). Anaerobic specimens were plated from frozen stocks within a Bactron II anaerobic chamber (Shel Lab, Cornelius, WA) using Brucella agar (Becton Dickinson, Franklin Lakes, NJ). For aerobic organisms, cultures were plated on trypticase soy agar with 5% sheep blood (Remel, Lenexa, KS) and incubated for 18 to 24 hr at 35°C. *Haemophilus influenzae* were grown on chocolate agar (Becton Dickinson, Franklin Lakes, NJ). *Streptococcus spp.* and *H. influenzae* were maintained at 5% CO_2_. *Campylobacter jejuni* were incubated at 40°C in microaerophilic conditions for 24-48 hours.

Following Clinical and Laboratory Standards Institute (CLSI) protocols, susceptibility testing was conducted using media specific to each organism (44–47). Aerobic isolates were tested in Cation-adjusted Mueller Hinton (CAMHB) broth (Becton Dickinson, Franklin Lakes, NJ). For *Streptococcus spp.*, *C. jejuni*, and *L. monocytogenes* CAMHB was supplemented with 3% laked horse blood (Hemostat, Dixon, CA). MIC testing of *H. influenzae* was performed in Haemophilus Test Medium Broth which was made by supplementing Mueller Hinton Broth (Becton Dickinson, Franklin Lakes, NJ) with 15 μg/mL nicotinamide adenine dinucleotide (Sigma-Aldrich, St. Louis, MO), 15 μg/mL porcine hematin (Sigma-Aldrich, St. Louis, MO), and 5 g/L yeast extract (Sigma-Aldrich, St. Louis, MO). Anaerobes were tested with Brucella broth (Becton Dickinson, Franklin Lakes, NJ) supplemented with 5 μg/mL hemin (Sigma-Aldrich, St. Louis, MO), 1 μg/mL Vitamin K1 (Sigma-Aldrich, St. Louis, MO), and 5% (v/v) LHB (Hemostat, Dixon, CA).

### Minimum inhibitory concentration assay

MICs were determined by broth microdilution performed by Microbiologics, Inc. (Kalamazoo, MI) according to CLSI guidelines (44, 46). Antibiotics were prepared as follows: aztreonam was dissolved in a saturated solution of sodium bicarbonate and diluted in water, chloramphenicol was dissolved in 95% ethanol and diluted in water, tedizolid was dissolved and diluted in DMSO, linezolid was dissolved and diluted in water, levofloxacin was dissolved in ½ volume of water then 0.1 mol/L NaOH dropwise and diluted in distilled water, and meropenem was dissolved and diluted in distilled water. Automated liquid handlers (Multidrop 384, Labsystems; Biomek 3000 and Biomek FX, Beckman Coulter) were used to perform two-fold serial dilutions. Plates were inoculated with a final bacterial concentration of 5 x 10^5^ CFU/mL. Aerobes were incubated aerobically at 35°C for 20 hours, *Campylobacter jejuni* were incubated at 40°C in a microaerophilic environment for 24 and 48 hours, and anaerobes were incubated anaerobically for 48 hours at 35°C. Following incubation, microplates were viewed using a plate viewer. For each of the test media and drugs, an un-inoculated solubility control plate was observed for evidence of drug precipitation in the test medium. No drug precipitation was observed. The MIC was recorded as the lowest concentration of drug that completely inhibited visible bacterial growth.

### Peptide synthesis

Peptide sequences are provided in table S11. A C-terminal cysteine was added to each peptide relative to its originally published sequence to facilitate conjugation to KLH or BSA. Peptides were synthesized using automated flow peptide synthesis and purified with reversed-phase high performance liquid chromatography by Amide Technologies (Cambridge, MA). The trifluoroacetate (TFA) counterion for the purified peptides was exchanged for hydrochloride. Peptides were >95% pure as verified using liquid chromatography-mass spectrometry on an Agilent G135B Single Quadrupole LC/MSD XT. Peptide amounts were confirmed via measurement at A205 before lyophilization. Peptides synthesized by Amide contained <1% residual TFA as measured via ionic chromatography by SB Peptide (Saint-Égrève, France). Additional L-Omp31-TB and D-Omp31-TB peptides for ELISAs were synthesized by GenScript (Piscataway, NJ) as TFA salts with >90% purity. Endotoxin content was determined using an Endosafe nexgen-PTS for all peptides and peptide conjugates. Peptides contained less than 0.05 EU/mL and peptide conjugates contained approximately 10-200 EU/mL.

### Peptide conjugation

Omp31-TB and OmpB_399_ were conjugated to KLH and StreptInCor was conjugated to BSA at particular peptide-to-carrier-protein ratios (Table S12) using a KLH- or BSA-peptide conjugation kit (CellMosaic, Woburn, MA). KLH conjugation with StreptInCor resulted in excessive precipitation that precluded buffer exchange and filtration and thus BSA was selected as the carrier protein for this peptide. Peptide labeling efficiency and loading was measured analytically via C18 HPLC and conjugate purity was determined via size-exclusion HPLC by CellMosaic. Peptide conjugate concentrations were quantified via a Pierce Bradford Protein Assay Kit (Thermo Fisher, Waltham, MA).

### Animals

Five- to six-week old female BALB/c mice (The Jackson Laboratory, Bar Harbor, ME) were maintained and handled at Avastus Preclinical Services (Cambridge, MA, USA). Mice were housed in groups of four to five for the immunization study or groups of three for the antibiotic toxicity study in Innovive disposable cages on an Innovive IVC mouse rack with free access to food (LabDiet 5053 PicoLab® Rodent Diet 20) and Innovive Aquavive® pre-filled water bottles. Mice were maintained on a twelve hour light/dark cycle at a temperature between 64°F and 72°F and 30–50% relative humidity. Experiments were only performed during the light phase. Mice were acclimated for one week prior to experiments and randomly distributed among the experimental groups. All animals were euthanized by CO_2_ asphyxiation at the end of all studies. Moribund animals or animals that lost 20% of their weight were euthanized. Animal experiments were conducted at Avastus Preclinical Services, Cambridge, MA, USA, under the standards set by its Institution Animal Care and Use Committee (IACUC), consistent with those of the Office of Laboratory Animal Welfare (OLAW), National Institutes of Health, USA. Avastus’ animal welfare assurance number is D16-00782 (A4543-01).

### Acute toxicity study

Mice were fasted for 3–4 hours prior to dosing. Tedizolid, linezolid, and chloramphenicol were ground into a fine powder using a mortar and pestle prior to resuspension in oral vehicle consisting of 0.5% wt/vol medium viscosity carboxymethylcellulose (Sigma-Aldrich, St. Louis, MO), 0.1% vol/vol Tween 80 (Fisher Scientific, Hampton, NH), and UltraPure distilled water (Thermo Fisher, Waltham, MA). Resuspended tedizolid, linezolid, and chloramphenicol were vortexed prior to dosing and administered at a fixed 200 µL volume via oral gavage with a 1 mL syringe and 20G disposable gavage needle. Aztreonam was completely dissolved in 43.8% wt/wt L-arginine (Thermo Fisher, Waltham, MA) and sterile 1X PBS. The aztreonam solution was sterilized via a 0.22 uM filter and administered in 200 µL intravenously via the tail vein. Two groups of control mice received either 200 µL of oral vehicle or intravenous vehicle (1X PBS alone). Food was reintroduced 1 hour after dosing all test articles. Mice were monitored for clinical signs 15–30 minutes, 1 hour, and 4 hours after dosing and daily thereafter. Animals were inspected for the following clinical signs: ataxia, body weight loss, dermatitis/skin lesion, diarrhea, hunched posture, labored breathing, lethargy, loss of coordination, low body temperature, piloerection, scruffed fur, self-isolation, seizure, shallow breathing, squinty eyes, tremors/shaking, ulcerated tumor, and weakness. Mice were weighed prior to dosing and daily thereafter.

### Hematological analysis

At the end of the acute toxicity study, blood was collected via cardiac puncture in K_2_EDTA tubes and stored at 4°C. The same day, a complete blood count was performed by IDEXX BioAnalytics (Grafton, MA) using a Sysmex XT-V automated hematology analyzer.

### Macroscopic examination and organ weights

At the end of the acute toxicity study, a full necropsy was performed and organ weights were recorded for the brain, ovaries, uterus, liver, kidneys, heart, spleen, lung, and thymus from all animals. Organs were also observed for lesions and abnormalities.

### Immunization

Mice were immunized once every 14 days 3–4 times depending on the immunogen (Fig 1A). Omp31-TB peptides and conjugates were administered in 30 µL intranasally (15 µL per nostril) alone or with 10 µg ODN 1826 VacciGrade (Invivogen, San Diego, CA). StreptInCor peptides and conjugates were administered in 100 µL subcutaneously across 1–2 sites. OmpB_399_ peptides and conjugates were administered in 100 µL subcutaneously for the initial dose and in 100 µL intraperitoneally thereafter. StreptInCor and OmpB_399_ peptides and conjugates were delivered alone or with Complete Freund’s adjuvant (1:1, v/v; Invivogen, San Diego, CA) for the first vaccination followed by Incomplete Freund’s Adjuvant thereafter (1:1, v/v; Invivogen, San Diego, CA). Three groups of mice were given PBS with the same volumes and routes of administration corresponding to each peptide group. Fourteen days after the final dose, blood was collected by cardiac puncture and processed to serum. BALF was collected using 1 mL of sterile PBS on day 56 for the Omp31-TB groups.

### ELISAs

Serum IgG and BALF IgA were quantified by ELISA. Peptides were diluted in 1 μg/mL Teknova Coating Buffer (Hollister, CA) and plated at 40 μL/well in 384-well MaxiSorp plates (Thermo Scientific, Waltham, MA) for overnight incubation at 4°C. Plates were washed with PBST (pH 7.4, 0.05% Tween-20), blocked with ChonBlock (Chondrex, Woodinville, WA) for 1 hour at room temperature, and then washed and replaced with 30 μL/well serial dilutions in ChonBlock of serum (3-fold, starting at 1:100) or BALF (2-fold, starting at 1:2) in duplicate and incubated for 2 hours at room temperature. After washing, 30 μL/well of 1:3000 dilutions of HRP-conjugated rabbit anti-mouse IgG (Fisher Scientific, Hampton, NH) or goat anti-mouse IgA (Fisher Scientific, Hampton, NH) were added and incubated for 1 hour at room temperature, then washed prior to incubation with TMB solution (Abcam, Cambridge, UK) 30 μL/well for 30 minutes and addition of 30 μL/well stop solution (Abcam, Cambridge, UK). Absorbance values were measured using a microplate reader (SpectraMax i3x) at 450 nm. Endpoint titer was defined as the serum dilution that yielded an absorbance value greater than 0.4.

### Statistical analysis

All statistical analyses were performed using Prism (version 10.3.1) (GraphPad Software, LLC, Boston, MA). One-way ANOVA with Tukey’s or Dunnett’s *post-hoc* tests was used to compare groups as indicated in figures and tables. For the acute toxicity study, data from the two separately treated oral vehicle control groups were combined during statistical analysis.

## SUPPLEMENTARY MATERIALS

**Table S1.** Source numbers, Gram classifications, and oxygen requirements for bacterial strains and isolates used in antimicrobial susceptibility studies. ATCC, American Type Culture Collection; MMX, Microbiologics repository.

| Organism | Gram classification | Oxygen requirement | Source number |
| --- | --- | --- | --- |
| *Bacteroides fragilis* | Gram-negative | Obligate anaerobic | ATCC 25285 |
| *Bacteroides thetaiotaomicron* | Gram-negative | Obligate anaerobic | ATCC 29741 |
| *Campylobacter jejuni* | Gram-negative | Microaerophilic | ATCC 49300 |
| *Clostridioides difficile* | Gram-positive | Obligate anaerobic | ATCC 700057 |
| *Enterobacter cloacae* | Gram-negative | Facultative anaerobic | ATCC 13047 |
| *Enterobacter cloacae* (ATM-S) | Gram-negative | Facultative anaerobic | MMX 1335 |
| *Enterococcus faecalis* | Gram-positive | Facultative anaerobic | ATCC 29212 |
| *Escherichia coli* | Gram-negative | Facultative anaerobic | ATCC 25922 |
| *Haemophilus influenzae* | Gram-negative | Facultative anaerobic | ATCC 49247 |
| *Klebsiella pneumoniae* | Gram-negative | Facultative anaerobic | MMX 1350 |
| *Listeria monocytogenes* | Gram-positive | Facultative anaerobic | ATCC 19115 |
| *Pseudomonas aeruginosa* | Gram-negative | Facultative anaerobic | ATCC 27853 |
| *Salmonella enterica* | Gram-negative | Facultative anaerobic | ATCC 14028 |
| *Serratia marcescens* | Gram-negative | Facultative anaerobic | ATCC 8100 |
| *Shigella sonnei* | Gram-negative | Facultative anaerobic | ATCC 25931 |
| *Staphylococcus aureus* | Gram-positive | Facultative anaerobic | ATCC 29213 |
| *Streptococcus pneumoniae* | Gram-positive | Facultative anaerobic | ATCC 49619 |
| *Streptococcus pyogenes* | Gram-positive | Facultative anaerobic | ATCC 19615 |

**Table S2.** Clinical signs and gross pathological findings at necropsy for mice receiving parent or *ent*-chloramphenicol, -linezolid, -tedizolid, and -aztreonam (Excel file attached).

**Table S3.**
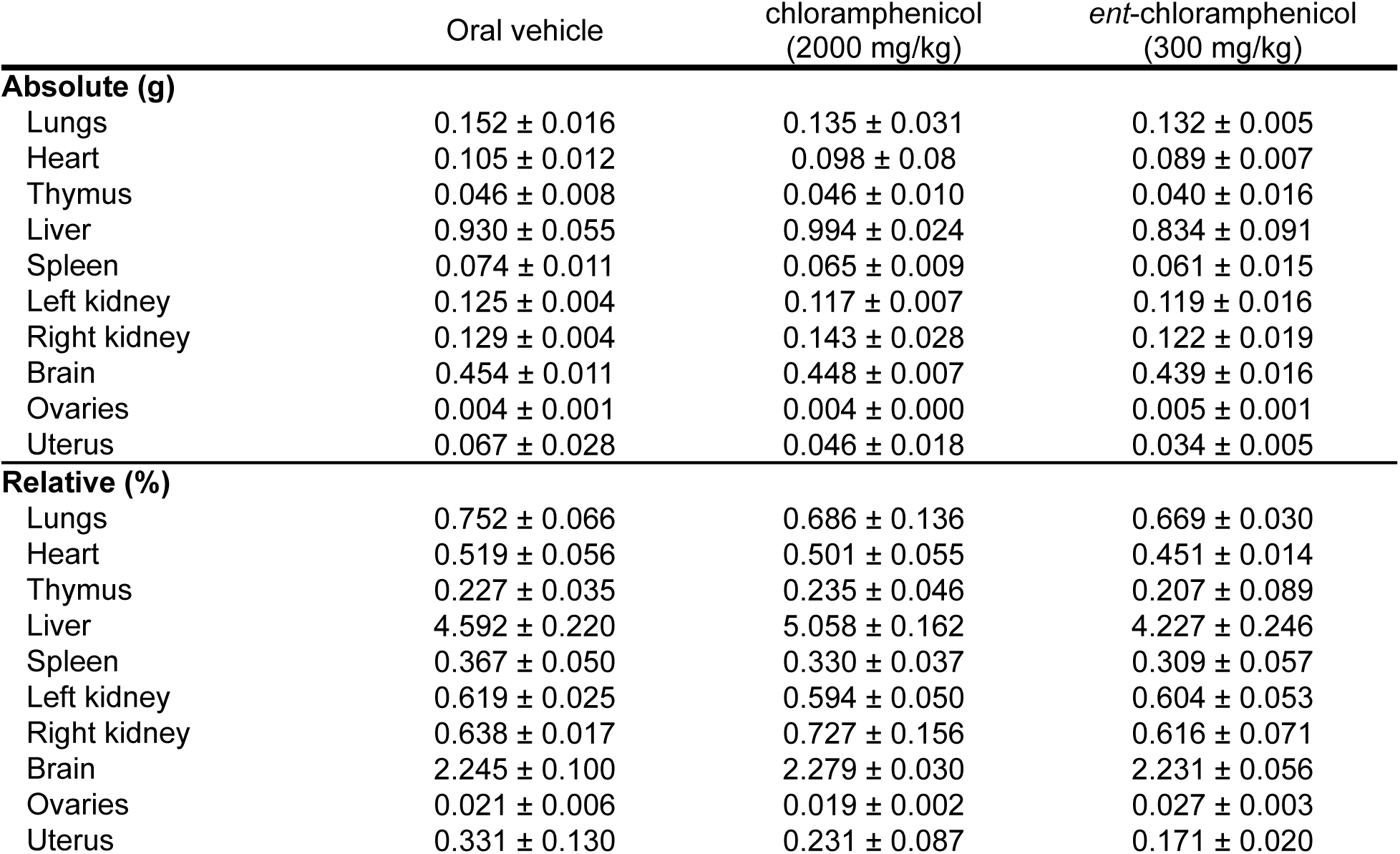
Absolute and relative organ weights (% of body weight) of animals treated with parent and *ent*-chloramphenicol on day 14. Results are expressed as mean ± SD. One-way ANOVA with Dunnett’s *post-hoc* tests was used to compare treatment groups with the control group. *, *p* ≤ 0.05; **, *p* ≤ 0.01; ***, *p* ≤ 0.001.

**Table S4.** Absolute and relative organ weights (% of body weight) of animals treated with parent and *ent*-linezolid on day 14. Results are expressed as mean ± SD. One-way ANOVA with Dunnett’s *post-hoc* tests was used to compare treatment groups with the control group. *, *p* ≤ 0.05; **, *p* ≤ 0.01; ***, *p* ≤ 0.001.

|  | Oral vehicle | linezolid<br>(2000 mg/kg) | ent-linezolid<br>(300 mg/kg) |
| --- | --- | --- | --- |
| **Absolute (g)** |  |  |  |
| Lungs | 0.152 ± 0.016 | 0.161 ± 0.016 | 0.156 ± 0.025 |
| Heart | 0.105 ± 0.012 | 0.099 ± 0.012 | 0.098 ± 0.004 |
| Thymus | 0.046 ± 0.008 | 0.039 ± 0.008 | 0.052 ± 0.001 |
| Liver | 0.930 ± 0.055 | 0.887 ± 0.058 | 0.922 ± 0.025 |
| Spleen | 0.074 ± 0.011 | 0.067 ± 0.007 | 0.062 ± 0.007 |
| Left kidney | 0.125 ± 0.004 | 0.127 ± 0.008 | 0.134 ± 0.007 |
| Right kidney | 0.129 ± 0.004 | 0.130 ± 0.022 | 0.134 ± 0.010 |
| Brain | 0.454 ± 0.011 | 0.438 ± 0.016 | 0.441 ± 0.013 |
| Ovaries | 0.004 ± 0.001 | 0.004 ± 0.001 | 0.006 ± 0.001 |
| Uterus | 0.067 ± 0.028 | 0.053 ± 0.037 | 0.051 ± 0.008 |
| **Relative (%)** |  |  |  |
| Lungs | 0.752 ± 0.066 | 0.773 ± 0.061 | 0.737 ± 0.128 |
| Heart | 0.519 ± 0.056 | 0.476 ± 0.051 | 0.461 ± 0.012 |
| Thymus | 0.227 ± 0.035 | 0.186 ± 0.040 | 0.247 ± 0.009 |
| Liver | 4.592 ± 0.220 | 4.271 ± 0.258 | 4.341 ± 0.139 |
| Spleen | 0.367 ± 0.050 | 0.324 ± 0.028 | 0.291 ± 0.033 |
| Left kidney | 0.619 ± 0.025 | 0.610 ± 0.035 | 0.632 ± 0.044 |
| Right kidney | 0.638 ± 0.017 | 0.628 ± 0.113 | 0.632 ± 0.058 |
| Brain | 2.245 ± 0.100 | 2.108 ± 0.042 | 2.062 ± 0.104 |
| Ovaries | 0.021 ± 0.006 | 0.017 ± 0.006 | 0.027 ± 0.005 |
| Uterus | 0.331 ± 0.130 | 0.254 ± 0.171 | 0.240 ± 0.035 |

**Table S5.** Absolute and relative organ weights (% of body weight) of animals treated with parent and *ent*-tedizolid on day 14. Results are expressed as mean ± SD. One-way ANOVA with Dunnett’s *post-hoc* tests was used to compare treatment groups with the control group. *, *p* ≤ 0.05; **, *p* ≤ 0.01; ***, *p* ≤ 0.001.

|  | Oral vehicle | tedizolid<br>(2000 mg/kg) | *ent*-tedizolid<br>(300 mg/kg) | *ent*-tedizolid<br>(2000 mg/kg) |
| --- | --- | --- | --- | --- |
| **Absolute (g)** |  |  |  |  |
| Lungs | 0.152 ± 0.016 | 0.140 ± 0.004 | 0.138 ± 0.011 | 0.129 ± 0.011 |
| Heart | 0.105 ± 0.012 | 0.097 ± 0.009 | 0.093 ± 0.009 | 0.095 ± 0.011 |
| Thymus | 0.046 ± 0.008 | 0.042 ± 0.005 | 0.047 ± 0.006 | 0.047 ± 0.005 |
| Liver | 0.930 ± 0.055 | 0.844 ± 0.041 | 0.857 ± 0.093 | 0.822 ± 0.099 |
| Spleen | 0.074 ± 0.011 | 0.072 ± 0.009 | 0.071 ± 0.007 | 0.078 ± 0.013 |
| Left kidney | 0.125 ± 0.004 | 0.129 ± 0.011 | 0.119 ± 0.012 | 0.127 ± 0.020 |
| Right kidney | 0.129 ± 0.004 | 0.141 ± 0.005 | 0.123 ± 0.015 | 0.119 ± 0.007 |
| Brain | 0.454 ± 0.011 | 0.441 ± 0.013 | 0.432 ± 0.012 | 0.431 ± 0.025 |
| Ovaries | 0.004 ± 0.001 | 0.003 ± 0.001 | 0.004 ± 0.001 | 0.004 ± 0.000 |
| Uterus | 0.067 ± 0.028 | 0.023 ± 0.005* | 0.029 ± 0.010 | 0.056 ± 0.007 |
| **Relative (%)** |  |  |  |  |
| Lungs | 0.752 ± 0.066 | 0.716 ± 0.005 | 0.676 ± 0.078 | 0.645 ± 0.029 |
| Heart | 0.519 ± 0.056 | 0.495 ± 0.038 | 0.452 ± 0.025 | 0.472 ± 0.039 |
| Thymus | 0.227 ± 0.035 | 0.215 ± 0.021 | 0.229 ± 0.023 | 0.237 ± 0.034 |
| Liver | 4.592 ± 0.220 | 4.316 ± 0.204 | 4.166 ± 0.294 | 4.091 ± 0.276* |
| Spleen | 0.367 ± 0.050 | 0.366 ± 0.044 | 0.347 ± 0.019 | 0.387 ± 0.039 |
| Left kidney | 0.619 ± 0.025 | 0.658 ± 0.062 | 0.579 ± 0.033 | 0.632 ± 0.052 |
| Right kidney | 0.638 ± 0.017 | 0.721 ± 0.017 | 0.598 ± 0.045 | 0.592 ± 0.011 |
| Brain | 2.245 ± 0.100 | 2.255 ± 0.125 | 2.104 ± 0.108 | 2.156 ± 0.170 |
| Ovaries | 0.021 ± 0.006 | 0.017 ± 0.003 | 0.019 ± 0.005 | 0.021 ± 0.002 |
| Uterus | 0.331 ± 0.130 | 0.117 ± 0.025* | 0.140 ± 0.043 | 0.283 ± 0.050 |

**Table S6.** Absolute and relative organ weights (% of body weight) of animals treated with parent and *ent*-aztreonam on day 14. Results are expressed as mean ± SD. One-way ANOVA with Dunnett’s *post-hoc* tests was used to compare treatment groups with the control group. *, *p* ≤ 0.05; **, *p* ≤ 0.01; ***, *p* ≤ 0.001.

|  | Intravenous vehicle | aztreonam<br>(1200 mg/kg) | ent-aztreonam<br>(400 mg/kg) | ent-aztreonam<br>(1200 mg/kg) |
| --- | --- | --- | --- | --- |
| **Absolute (g)** |  |  |  |  |
| Lungs | 0.139 ± 0.012 | 0.147 ± 0.010 | 0.174 ± 0.021 | 0.166 ± 0.032 |
| Heart | 0.101 ± 0.012 | 0.106 ± 0.009 | 0.117 ± 0.014 | 0.113 ± 0.011 |
| Thymus | 0.050 ± 0.004 | 0.050 ± 0.004 | 0.068 ± 0.007* | 0.058 ± 0.011 |
| Liver | 0.958 ± 0.106 | 1.021 ± 0.086 | 1.082 ± 0.124 | 1.064 ± 0.125 |
| Spleen | 0.061 ± 0.010 | 0.068 ± 0.006 | 0.069 ± 0.008 | 0.068 ± 0.009 |
| Left kidney | 0.130 ± 0.016 | 0.135 ± 0.010 | 0.140 ± 0.008 | 0.133 ± 0.019 |
| Right kidney | 0.135 ± 0.012 | 0.143 ± 0.013 | 0.151 ± 0.012 | 0.135 ± 0.014 |
| Brain | 0.430 ± 0.020 | 0.451 ± 0.012 | 0.469 ± 0.012 | 0.451 ± 0.022 |
| Ovaries | 0.005 ± 0.000 | 0.005 ± 0.001 | 0.005 ± 0.000 | 0.005 ± 0.002 |
| Uterus | 0.094 ± 0.003 | 0.134 ± 0.038 | 0.128 ± 0.029 | 0.124 ± 0.034 |
| **Relative (%)** |  |  |  |  |
| Lungs | 0.719 ± 0.037 | 0.734 ± 0.037 | 0.807 ± 0.062 | 0.806 ± 0.205 |
| Heart | 0.520 ± 0.012 | 0.530 ± 0.067 | 0.547 ± 0.091 | 0.544 ± 0.076 |
| Thymus | 0.259 ± 0.023 | 0.253 ± 0.041 | 0.315 ± 0.041 | 0.277 ± 0.049 |
| Liver | 4.955 ± 0.109 | 5.075 ± 0.223 | 5.007 ± 0.347 | 5.102 ± 0.273 |
| Spleen | 0.313 ± 0.022 | 0.337 ± 0.013 | 0.321 ± 0.021 | 0.326 ± 0.047 |
| Left kidney | 0.674 ± 0.012 | 0.673 ± 0.027 | 0.649 ± 0.037 | 0.638 ± 0.037 |
| Right kidney | 0.701 ± 0.051 | 0.710 ± 0.017 | 0.698 ± 0.047 | 0.647 ± 0.029 |
| Brain | 2.240 ± 0.208 | 2.246 ± 0.145 | 2.177 ± 0.062 | 2.172 ± 0.132 |
| Ovaries | 0.026 ± 0.003 | 0.026 ± 0.003 | 0.023 ± 0.000 | 0.024 ± 0.008 |
| Uterus | 0.492 ± 0.078 | 0.670 ± 0.184 | 0.594 ± 0.132 | 0.602 ± 0.196 |

**Table S7.** Hematological parameters of animals treated with parent and *ent*-chloramphenicol on day 14. Results are expressed as mean ± SD. One-way ANOVA with Dunnett’s *post-hoc* tests was used to compare treatment groups with the control group. *, *p* ≤ 0.05; **, *p* ≤ 0.01; ***, *p* ≤ 0.001. WBC, white blood cells; RBC, red blood cells; HGB, hemoglobin; HCT, hematocrit; MCV, mean corpuscular volume; MCH, mean corpuscular hemoglobin; MCHC, mean corpuscular hemoglobin concentration; PLT, platelets.

|  | Oral vehicle | chloramphenicol<br>(2000 mg/kg) | ent-chloramphenicol<br>(300 mg/kg) |
| --- | --- | --- | --- |
| WBC ($10^3/\mu\text{L}$) | 8.35 ± 3.56 | 7.30 ± 2.43 | 5.83 ± 1.83 |
| RBC ($10^6/\mu\text{L}$) | 10.03 ± 0.39 | 10.19 ± 0.50 | 10.37 ± 0.22 |
| HGB (g/dL) | 15.08 ± 0.53 | 15.50 ± 0.72 | 15.40 ± 0.30 |
| HCT (%) | 48.0 ± 1.5 | 48.6 ± 2.1 | 49.4 ± 1.1 |
| MCV (fL) | 47.7 ± 0.5 | 48.0 ± 0.0 | 48.0 ± 0.0 |
| MCH (pg) | 15.05 ± 0.12 | 15.20 ± 0.10 | 14.83 ± 0.06 |
| MCHC (g/dL) | 31.4 ± 0.3 | 31.9 ± 0.3 | 31.2 ± 0.2 |
| PLT ($10^3/\mu\text{L}$) | 969 ± 227 | 1123 ± 83 | 655 ± 291 |
| Neutrophils (/$\mu\text{L}$) | 935 ± 385 | 913 ± 613 | 827 ± 237 |
| Lymphocytes (/$\mu\text{L}$) | 6862 ± 2997 | 6026 ± 1965 | 4688 ± 1786 |
| Monocytes (/$\mu\text{L}$) | 391 ± 246 | 192 ± 66 | 278 ± 146 |
| Eosinophils (/$\mu\text{L}$) | 153 ± 75 | 161 ± 222 | 41 ± 40 |
| Basophils (/$\mu\text{L}$) | 9.00 ± 8.92 | 8.00 ± 6.93 | 0.00 ± 0.00 |
| Reticulocytes ($10^3/\mu\text{L}$) | 351 ± 47 | 312 ± 20 | 266 ± 37 |

**Table S8.** Hematological parameters of animals treated with parent and *ent*-linezolid on day 14. Results are expressed as mean ± SD. One-way ANOVA with Dunnett’s *post-hoc* tests was used to compare treatment groups with the control group. *, *p* ≤ 0.05; **, *p* ≤ 0.01; ***, *p* ≤ 0.001. WBC, white blood cells; RBC, red blood cells; HGB, hemoglobin; HCT, hematocrit; MCV, mean corpuscular volume; MCH, mean corpuscular hemoglobin; MCHC, mean corpuscular hemoglobin concentration; PLT, platelets.

|  | Oral vehicle | linezolid<br>(2000 mg/kg) | ent-linezolid<br>(300 mg/kg) |
| --- | --- | --- | --- |
| WBC ($10^3/\mu\text{L}$) | 8.35 ± 3.56 | 8.37 ± 1.57 | 7.77 ± 5.46 |
| RBC ($10^6/\mu\text{L}$) | 10.03 ± 0.39 | 9.90 ± 0.24 | 10.30 ± 0.22 |
| HGB (g/dL) | 15.08 ± 0.53 | 15.17 ± 0.49 | 15.50 ± 0.27 |
| HCT (%) | 48.0 ± 1.5 | 47.7 ± 0.7 | 49.3 ± 0.5 |
| MCV (fL) | 47.7 ± 0.5 | 48.3 ± 0.6 | 48.0 ± 1.0 |
| MCH (pg) | 15.05 ± 0.12 | 15.33 ± 0.12 | 15.03 ± 0.06 |
| MCHC (g/dL) | 31.4 ± 0.3 | 31.8 ± 0.7 | 31.4 ± 0.3 |
| PLT ($10^3/\mu\text{L}$) | 969 ± 227 | 878 ± 128 | 1069 ± 333 |
| Neutrophils (/$\mu\text{L}$) | 935 ± 385 | 973 ± 82 | 722 ± 398 |
| Lymphocytes (/$\mu\text{L}$) | 6862 ± 2997 | 6967 ± 1545 | 6567 ± 4782 |
| Monocytes (/$\mu\text{L}$) | 391 ± 246 | 312 ± 156 | 326 ± 183 |
| Eosinophils (/$\mu\text{L}$) | 153 ± 75 | 111 ± 30 | 144 ± 125 |
| Basophils (/$\mu\text{L}$) | 9.00 ± 8.92 | 3.33 ± 5.77 | 7.00 ± 6.25 |
| Reticulocytes ($10^3/\mu\text{L}$) | 351 ± 47 | 413 ± 57 | 288 ± 41 |

**Table S9.**
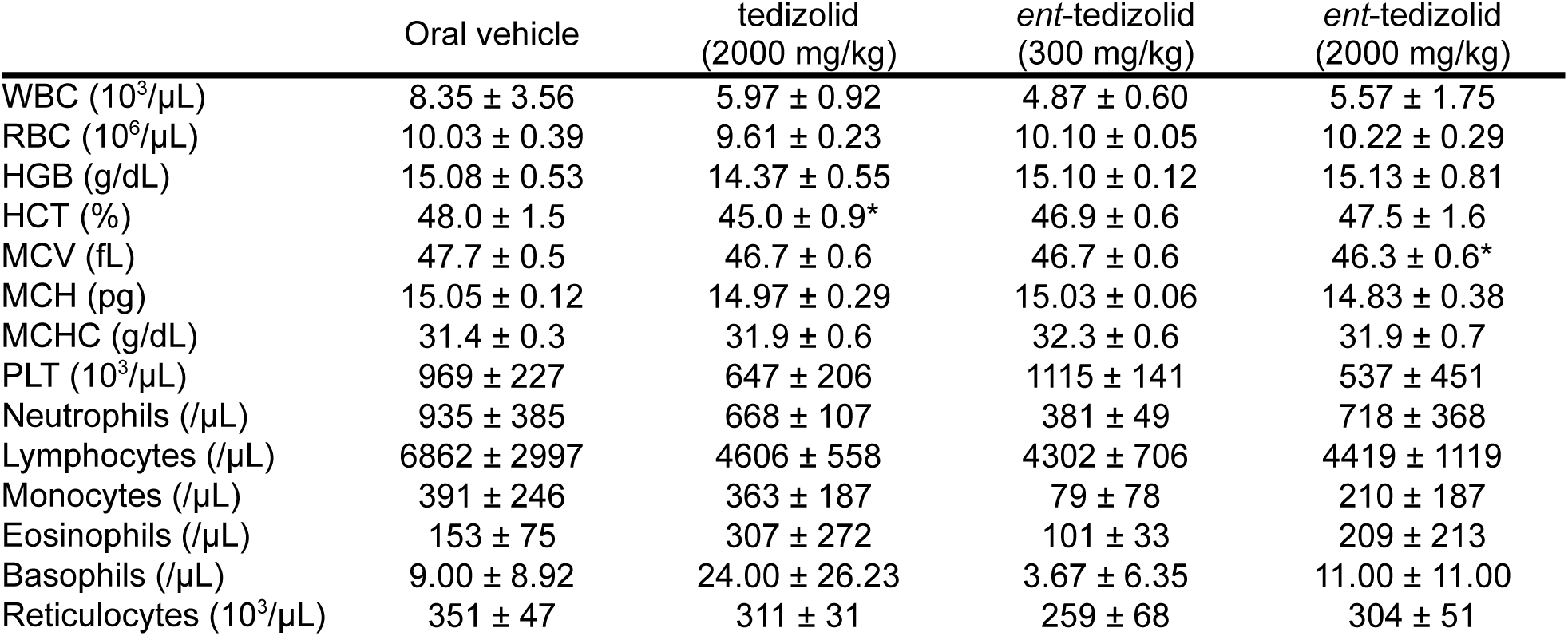
Hematological parameters of animals treated with parent and *ent*-tedizolid on day 14. Results are expressed as mean ± SD. One-way ANOVA with Dunnett’s *post-hoc* tests was used to compare treatment groups with the control group. *, *p* ≤ 0.05; **, *p* ≤ 0.01; ***, *p* ≤ 0.001. WBC, white blood cells; RBC, red blood cells; HGB, hemoglobin; HCT, hematocrit; MCV, mean corpuscular volume; MCH, mean corpuscular hemoglobin; MCHC, mean corpuscular hemoglobin concentration; PLT, platelets.

**Table S10.** Hematological parameters of animals treated with parent and *ent*-aztreonam on day 14. Results are expressed as mean ± SD. One-way ANOVA with Dunnett’s *post-hoc* tests was used to compare treatment groups with the control group. *, *p* ≤ 0.05; **, *p* ≤ 0.01; ***, *p* ≤ 0.001. WBC, white blood cells; RBC, red blood cells; HGB, hemoglobin; HCT, hematocrit; MCV, mean corpuscular volume; MCH, mean corpuscular hemoglobin; MCHC, mean corpuscular hemoglobin concentration; PLT, platelets.

|  | Intravenous vehicle | aztreonam<br>(1200 mg/kg) | ent-aztreonam<br>(400 mg/kg) | ent-aztreonam<br>(1200 mg/kg) |
| --- | --- | --- | --- | --- |
| WBC (10<sup>3</sup>/μL) | 10.00 ± 2.41 | 5.97 ± 1.90 | 11.03 ± 1.34 | 9.40 ± 2.81 |
| RBC (10<sup>6</sup>/μL) | 10.31 ± 0.16 | 10.28 ± 0.34 | 10.18 ± 0.37 | 10.20 ± 0.40 |
| HGB (g/dL) | 15.70 ± 0.10 | 15.50 ± 0.46 | 15.37 ± 0.38 | 15.4 ± 0.46 |
| HCT (%) | 50.7 ± 0.4 | 49.8 ± 1.1 | 49.8 ± 1.0 | 49.6 ± 1.4 |
| MCV (fL) | 49.3 ± 0.6 | 48.7 ± 0.6 | 48.7 ± 0.6 | 48.3 ± 0.6 |
| MCH (pg) | 15.23 ± 0.15 | 15.07 ± 0.15 | 15.10 ± 0.20 | 15.10 ± 0.27 |
| MCHC (g/dL) | 30.9 ± 0.1 | 31.1 ± 0.4 | 30.9 ± 0.2 | 31.1 ± 0.3 |
| PLT (10<sup>3</sup>/μL) | 833 ± 49 | 865 ± 52 | 870 ± 47 | 826 ± 89 |
| Neutrophils (/μL) | 823 ± 161 | 886 ± 180 | 1274 ± 233 | 962 ± 299 |
| Lymphocytes (/μL) | 8504 ± 2342 | 4692 ± 1647 | 9148 ± 1286 | 7977 ± 2575 |
| Monocytes (/μL) | 550 ± 38 | 280 ± 45 | 477 ± 93 | 364 ± 236 |
| Eosinophils (/μL) | 114 ± 29 | 95 ± 37 | 124 ± 8 | 90 ± 40 |
| Basophils (/μL) | 10.00 ± 2.00 | 14.33 ± 7.77 | 11.33 ± 1.16 | 7.00 ± 6.25 |
| Reticulocytes (10<sup>3</sup>/μL) | 405 ± 103 | 360 ± 20 | 346 ± 6 | 356 ± 24 |

**Table S11.**
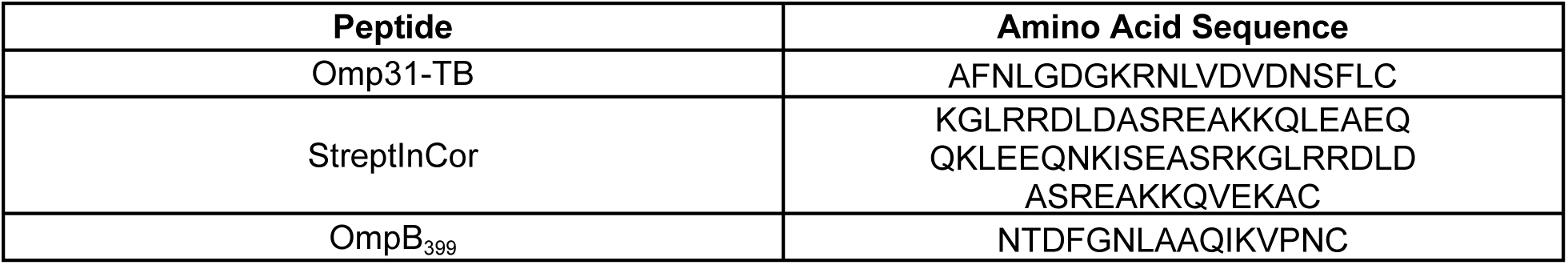
Amino acid sequences of peptides used in this study. A C-terminal cysteine was added to facilitate conjugation to KLH and BSA.

**Table S12.** Peptide loading per carrier protein.

| Peptide Conjugate | Number of Peptides per Carrier |
| --- | --- |
| L-Omp31-TB KLH | 13.78 |
| D-Omp31-TB KLH | 10.31 |
| L-StreptInCor BSA | 0.3 |
| D-StreptInCor BSA | 0.4 |
| L-OmpB<sub>399</sub> KLH | 13.8 |
| D-OmpB<sub>399</sub> KLH | 18 |

**Figure S1.**
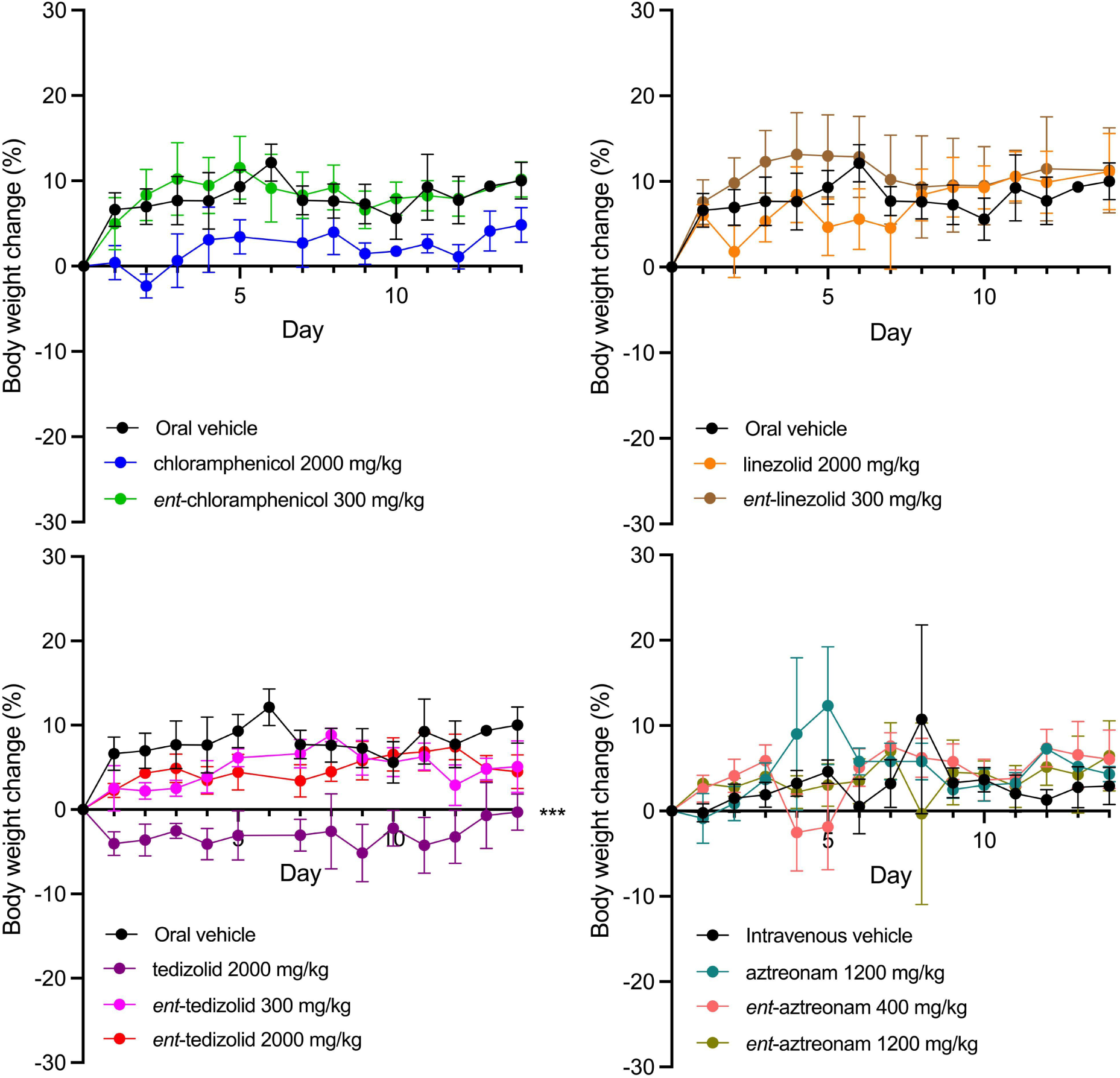
Body weight changes of mice over time after treatment with parent or *ent*-chloramphenicol, -linezolid, -tedizolid, and -aztreonam. Data is plotted as mean ± SD. One-way ANOVA with Dunnett’s *post-hoc* tests was used to compare treatment groups with their respective control groups on day 14. *, *p* ≤ 0.05; **, *p* ≤ 0.01; ***, *p* ≤ 0.001.

## REFERENCES

1. Adamala KP, Agashe D, Belkaid Y, Bittencourt DM de C, Cai Y, Chang MW, Chen IA, Church GM, Cooper VS, Davis MM, Devaraj NK, Endy D, Esvelt KM, Glass JI, Hand TW, Inglesby TV, Isaacs FJ, James WG, Jones JDG, Kay MS, Lenski RE, Liu C, Medzhitov R, Nicotra ML, Oehm SB, Pannu J, Relman DA, Schwille P, Smith JA, Suga H, Szostak JW, Talbot NJ, Tiedje JM, Venter JC, Winter G, Zhang W, Zhu X, Zuber MT. 2024. Confronting risks of mirror life. Science 0:eads9158.

2. Blackmond DG. 2010. The Origin of Biological Homochirality. Cold Spring Harb Perspect Biol 2:a002147.

3. Adamala KP, Esvelt KM, Glass JI, Oehm SB, Suga H, Szostak JW. 11 December 2024, posting date. Chapter 2: Pathways to Mirror Life. In Technical Report on Mirror Bacteria: Feasibility and Risks. Stanford Digital Repository, Stanford, California. doi:10.25740/cv716pj4036.

4. Agashe D, Binder DJ, Cooper VS, Esvelt KM, Lenski RE, Relman DA. 11 December 2024, posting date. Chapter 8: Environmental Survival and Spread. In Technical Report on Mirror Bacteria: Feasibility and Risks. Stanford Digital Repository, Stanford, California. doi:10.25740/cv716pj4036.

5. Duncombe RK, Hand TW, Inglesby TV, Lewis G, Medzhitov R, Nicotra ML, Pannu J, Relman DA, Sweere JM. 11 December 2024, posting date. Chapter 4: Risks to Human Health. In Technical Report on Mirror Bacteria: Feasibility and Risks. Stanford Digital Repository, Stanford, California. doi:10.25740/cv716pj4036.

6. Inglesby TV, Lewis G, Relman DA, Sweere JM, Wang B. 11 December 2024, posting date. Chapter 5: Medical Countermeasures. In Technical Report on Mirror Bacteria: Feasibility and Risks. Stanford Digital Repository, Stanford, California. doi:10.25740/cv716pj4036.

7. Jones LH. 2015. Recent advances in the molecular design of synthetic vaccines. Nat Chem 7:952–960.

8. Rappuoli R, De Gregorio E, Costantino P. 2019. On the mechanisms of conjugate vaccines. Proc Natl Acad Sci 116:14–16.

9. Malonis RJ, Lai JR, Vergnolle O. 2020. Peptide-Based Vaccines: Current Progress and Future Challenges. Chem Rev 120:3210–3229.

10. Hayakawa I, Atarashi S, Yokohama S, Imamura M, Sakano K, Furukawa M. 1986. Synthesis and antibacterial activities of optically active ofloxacin. Antimicrob Agents Chemother 29:163–164.

11. Culbertson TP, Domagala JM, Nichols JB, Priebe S, Skeean RW. 1987. Enantiomers of 1-ethyl-7-[3-[(ethylamino)methyl]-1-pyrrolidinyl]-6,8-difluoro-1,4-dihydro-4-oxo-3-qui nolinecarboxylic acid: preparation and biological activity. J Med Chem 30:1711–1715.

12. Pedroni L, Dall’Asta C, Galaverna G, Dellafiora L. 2025. Computational perspectives on amoxicillin and Staphylococcus aureus in mirror life. Glob Chall 9:e00051.

13. Muller S, Guichard G, Benkirane N, Brown F, Van Regenmortel MH, Briand JP. 1995. Enhanced immunogenicity and cross-reactivity of retro-inverso peptidomimetics of the major antigenic site of foot-and-mouth disease virus. Pept Res 8:138–144.

14. Briand J-P, Benkirane N, Guichard G, Newman JFE, Van Regenmortel MHV, Brown F, Muller S. 1997. A retro-inverso peptide corresponding to the GH loop of foot-and-mouth disease virus elicits high levels of long-lasting protective neutralizing antibodies. Proc Natl Acad Sci 94:12545–12550.

15. Ben-Yedidia T, Beignon A-S, Partidos CD, Muller S, Arnon R. 2002. A retro-inverso peptide analogue of influenza virus hemagglutinin B-cell epitope 91-108 induces a strong mucosal and systemic immune response and confers protection in mice after intranasal immunization. Mol Immunol 39:323–331.

16. Fromme B, Eftekhari P, Van Regenmortel M, Hoebeke J, Katz A, Millar R. 2003. A novel retro-inverso gonadotropin-releasing hormone (GnRH) immunogen elicits antibodies that neutralize the activity of native GnRH. Endocrinology 144:3262–3269.

17. Fischer P, Comis A, Tyler M, Howden M. 2007. Oral and parenteral immunization with synthetic retro-inverso peptides induce antibodies that cross-react with native peptides and parent antigens. Indian J Biochem Biophys 44:140–144.

18. Chong P, Sia C, Tripet B, James O, Klein M. 1996. Comparative immunological properties of enantiomeric peptides. Lett Pept Sci 3:99–106.

19. Zanichelli V, Sharland M, Cappello B, Moja L, Getahun H, Pessoa-Silva C, Sati H, van Weezenbeek C, Balkhy H, Simão M, Gandra S, Huttner B. 2023. The WHO AWaRe (Access, Watch, Reserve) antibiotic book and prevention of antimicrobial resistance. Bull World Health Organ 101:290–296.

20. M100 Ed34 | Performance Standards for Antimicrobial Susceptibility Testing, 34th Edition. Clin Lab Stand Inst. https://clsi.org/standards/products/microbiology/documents/m100/. Retrieved 3 December 2024.

21. Glass AM, Coombs W, Taffet SM. 2013. Spontaneous Cardiac Calcinosis in BALB/cByJ Mice. Comp Med 63:29–37.

22. Wang P, Xiong X, Jiao J, Yang X, Jiang Y, Wen B, Gong W. 2017. Th1 epitope peptides induce protective immunity against *Rickettsia rickettsii* infection in C3H/HeN mice. Vaccine 35:7204–7212.

23. Postol E, Alencar R, Higa FT, Barros SF de, Demarchi LMF, Kalil J, Guilherme L. 2013. StreptInCor: A Candidate Vaccine Epitope against S. pyogenes Infections Induces Protection in Outbred Mice. PLOS ONE 8:e60969.

24. Zhang F, Li Z, Jia B, Zhu Y, Pang P, Zhang C, Ding J. 2019. The Immunogenicity of OMP31 Peptides and Its Protection Against Brucella melitensis Infection in Mice. Sci Rep 9:3512.

25. Hervé M, Maillére B, Mourier G, Texier C, Leroy S, Ménez A. 1997. On the immunogenic properties of retro-inverso peptides. Total retro-inversion of T-Cell epitopes causes a loss of binding to MHC II molecules. Mol Immunol 34:157–163.

26. Purcell AW, McCluskey J, Rossjohn J. 2007. More than one reason to rethink the use of peptides in vaccine design. Nat Rev Drug Discov 6:404–414.

27. Black M, Trent A, Tirrell M, Olive C. 2010. Advances in the design and delivery of peptide subunit vaccines with a focus on Toll-like receptor agonists. Expert Rev Vaccines 9:157–173.

28. Maxwell RE, Nickel VS. 1954. The antibacterial activity of the isomers of chloramphenicol. Antibiot Chemother Northfield Ill 4:289–295.

29. Brock TD. 1961. CHLORAMPHENICOL. Bacteriol Rev 25:32–48.

30. Gregory WA, Brittelli DR, Wang CLJ, Wuonola MA, McRipley RJ, Eustice DC, Eberly VS, Slee AM, Forbes M, Bartholomew PT. 1989. Antibacterials. Synthesis and structure-activity studies of 3-aryl-2-oxooxazolidines. 1. The B group. J Med Chem 32:1673–1681.

31. Barbachyn MR, Ford CW. 2003. Oxazolidinone structure-activity relationships leading to linezolid. Angew Chem Int Ed Engl 42:2010–2023.

32. Fortuna CG, Berardozzi R, Bonaccorso C, Caltabiano G, Di Bari L, Goracci L, Guarcello A, Pace A, Palumbo Piccionello A, Pescitelli G, Pierro P, Lonati E, Bulbarelli A, Cocuzza CEA, Musumarra G, Musumeci R. 2014. New potent antibacterials against Gram-positive multiresistant pathogens: Effects of side chain modification and chirality in linezolid-like 1,2,4-oxadiazoles. Bioorg Med Chem 22:6814–6825.

33. Breuer H, Cimarusti CM, Denzel Th, Koster WH, Slusarchyk WA, Treuner UD. 1981. Monobactams—structure-activity relationships leading to SQ 26,776. J Antimicrob Chemother 8:21–28.

34. Decuyper L, Jukič M, Sosič I, Žula A, D’hooghe M, Gobec S. 2018. Antibacterial and β-Lactamase Inhibitory Activity of Monocyclic β-Lactams. Med Res Rev 38:426–503.

35. Tatsuta K. 2013. Total synthesis of the big four antibiotics and related antibiotics. J Antibiot (Tokyo) 66:107–129.

36. 2020. M3(R2) Nonclinical Safety Studies for the Conduct of Human Clinical Trials and Marketing Authorization for Pharmaceuticals. FDA. https://www.fda.gov/regulatory-information/search-fda-guidance-documents/m3r2-nonclinical-safety-studies-conduct-human-clinical-trials-and-marketing-authorization. Retrieved 3 December 2024.

37. 2009. ICH M3 (R2) Non-clinical safety studies for the conduct of human clinical trials for pharmaceuticals - Scientific guideline | European Medicines Agency (EMA). https://www.ema.europa.eu/en/ich-m3-r2-non-clinical-safety-studies-conduct-human-clinical-trials-pharmaceuticals-scientific-guideline. Retrieved 16 December 2024.

38. Muller S, Benkirane N, Guichard G, Van Regenmortel MH, Brown F. 1998. The potential of retro-inverso peptides as synthetic vaccines. Expert Opin Investig Drugs 7:1429–1438.

39. Sharon J, Rynkiewicz MJ, Lu Z, Yang C-Y. 2014. Discovery of protective B-cell epitopes for development of antimicrobial vaccines and antibody therapeutics. Immunology 142:1–23.

40. Plotkin SA. 2010. Correlates of Protection Induced by Vaccination. Clin Vaccine Immunol 17:1055–1065.

41. Motley MP, Diago-Navarro E, Banerjee K, Inzerillo S, Fries BC. 2020. The Role of IgG Subclass in Antibody-Mediated Protection against Carbapenem-Resistant Klebsiella pneumoniae. mBio 11:10.1128/mbio.02059-20.

42. Taylor JM, Ziman ME, Canfield DR, Vajdy M, Solnick JV. 2008. Effects of a Th1 versus a Th2 Biased Immune Response in Protection Against Helicobacter pylori Challenge in Mice. Microb Pathog 44:20–27.

43. Rappuoli R. 2018. Glycoconjugate vaccines: Principles and mechanisms. Sci Transl Med 10:eaat4615.

44. M11 Ed9 Anaerobic Bacteria Antimicrobial Susceptibility. Clin Lab Stand Inst. https://clsi.org/standards/products/microbiology/documents/m11/. Retrieved 10 December 2024.

45. M02 Ed14 | Performance Standards for Antimicrobial Disk Susceptibility Tests, 14th Edition. Clin Lab Stand Inst. https://clsi.org/standards/products/microbiology/documents/m02/. Retrieved 10 December 2024.

46. M07 Ed12 | Methods for Dilution Antimicrobial Susceptibility Tests for Bacteria That Grow Aerobically, 12th Edition. Clin Lab Stand Inst. https://clsi.org/standards/products/microbiology/documents/m07/. Retrieved 10 December 2024.

47. M45 Ed3 Test Infrequently Isolated/Fastidious Bacteria. Clin Lab Stand Inst. https://clsi.org/standards/products/microbiology/documents/m45/. Retrieved 10 December 2024.

